# AmpliconSuite: an end-to-end workflow for analyzing focal amplifications in cancer genomes

**DOI:** 10.1101/2024.05.06.592768

**Authors:** Jens Luebeck, Edwin Huang, Forrest Kim, Ted Liefeld, Bhargavi Dameracharla, Rohil Ahuja, Daniel Schreyer, Gino Prasad, Michał Adamaszek, Rishaan Kenkre, Tushar Agashe, Devika Torvi, Thorin Tabor, Mădălina Giurgiu, Soyeon Kim, Hoon Kim, Peter Bailey, Roel G.W. Verhaak, Viraj Deshpande, Michael Reich, Paul S. Mischel, Jill Mesirov, Vineet Bafna

## Abstract

Focal amplifications in the cancer genome, particularly extrachromosomal DNA (ecDNA) amplifications, are emerging as a pivotal event in cancer progression across diverse cancer contexts, presenting a paradigm shift in our understanding of tumor dynamics. Simultaneously, identification of the various modes of focal amplifications is bioinformatically challenging. We present AmpliconSuite, a collection of tools that enables robust identification of focal amplifications from whole-genome sequencing data. AmpliconSuite includes AmpliconSuite-pipeline; utilizing the AmpliconArchitect (AA) method, and AmpliconRepository.org; a community-editable website for the sharing of focal amplification calls. We also describe improvements made to AA since its initial release that improve its accuracy and speed. As a proof of principle, we utilized publicly available pan-cancer datasets encompassing 2,525 tumor samples hosted on AmpliconRepository.org to illustrate important properties of focal amplifications, showing ecDNA has higher copy number, and stronger oncogene enrichment, compared to other classes of focal amplifications. Finally, we illustrate how AmpliconSuite-pipeline enables delineation of the various mechanisms by which ecDNA forms.

## INTRODUCTION

The focal amplification of oncogenes is a frequent driver of cancer development, yet different modes of amplification have both distinct cellular and clinical consequences. For example, extrachromosomal DNA (ecDNA)-mediated amplifications are associated with high transcription levels^1–3^, rapid copy-number adaptation and therapeutic resistance^4–7^. Distinguishing different modes of focal amplification using whole genome sequencing (WGS) data is bioinformatically challenging^8^, and there is a need for a robust toolset to discover these events, as well as a need for a community-driven database to share the resulting focal amplification call sets.

We present AmpliconSuite and its companion web platform, AmpliconRepository.org, which offer a standardized workflow for focal amplification analysis and a community-sourced platform for result sharing. At the core of AmpliconSuite is the AmpliconArchitect (AA) method^9^. AA jointly analyzes structural variants (SVs) and copy numbers (CNs) from WGS data, and has been deployed on thousands of samples^10–15^, to elucidate the critical role of ecDNA in cancer. To make the AA method more easily accessible to the broadest spectrum of cancer investigators, we created AmpliconSuite-pipeline. AmpliconSuite-pipeline provides an end-to-end wrapper for AA and its associated focal amplification classification tool AmpliconClassifier (AC). Importantly, AmpliconSuite-pipeline introduces a methodology for preparing standardized inputs to AA, improving the reproducibility and interpretability of outputs, which functions on both human and mouse genome data. We deployed this workflow to GenePattern^16^ and Nextflow^17^ interfaces, enabling usage without bioinformatic tool installation. Simultaneously, AmpliconRepository.org enables community-sourced sharing of the focal amplification calls produced by AmpliconSuite. Here we describe AmpliconSuite and AmpliconRepository, and their application to a large collection of publicly available cancer samples, and describe enhancements implemented in AA since its initial release.

## RESULTS

### AmpliconSuite-pipeline

AmpliconSuite-pipeline provides robust identification of focal amplifications while also enabling users to enter the workflow from any intermediate point. If a user begins with FASTQ files, AmpliconSuite-pipeline performs alignment to a reference using a combination of BWA-MEM^18^ and samtools^19^ (Figure 1). With the resulting BAM file, or the user’s self-provided BAM or CRAM file, AmpliconSuite-pipeline next performs identification of candidate regions of focal amplification from whole-genome copy number calls prior to launching AmpliconArchitect. Unless users provide a BED file of CNV calls, AmpliconSuite-pipeline will invoke CNVkit^20^ for this task. Those CNV calls are not propagated into AA, rather they are used to establish candidate focal amplification locations, called “seed regions,” where AA launches its search. AmpliconSuite-pipeline implements a routine for removing regions from the CNV calls not consistent with focal amplifications (Methods – Seed selection). In brief, AmpliconSuite-pipeline identifies seed regions by filtering regions not more than two copies above median chromosome arm ploidy. Filters are also applied based on amplification size and include previously published^9^ AA filters (*amplified_intervals.py*) for repetitive and poorly mappable sequences along regions of the reference genome commonly observed to have elevated copy number in non-cancer samples.

**Figure 1:**
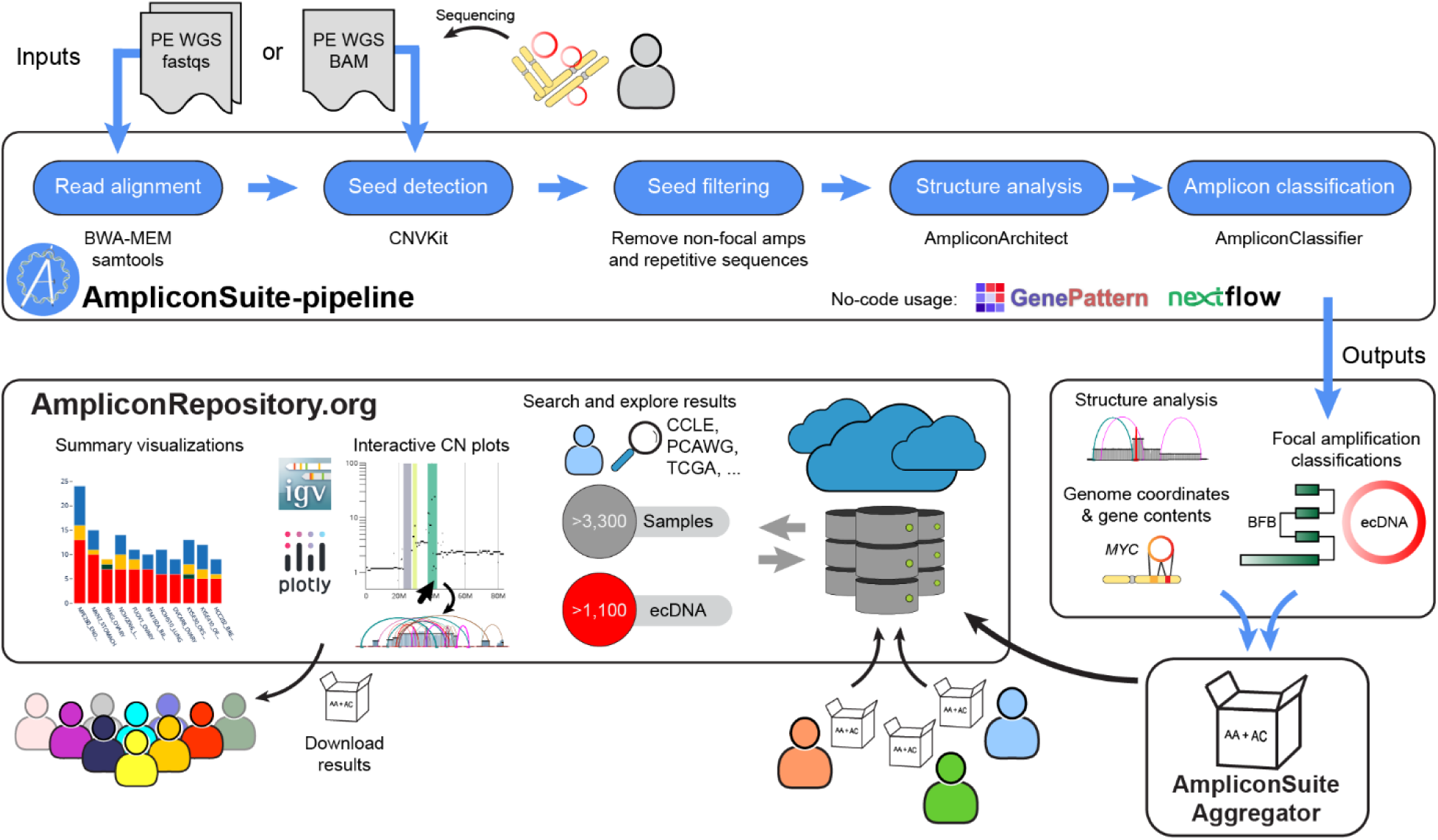
Schematic of AmpliconSuite-pipeline and AmpliconRepository. A cartoon schematic showing AmpliconSuite-pipeline and AmpliconRepository. Users provide sequencing data to AmpliconSuite-pipeline, which invokes alignment and genome-wide CNV calling for the purposes of identifying candidate regions of focal amplification. Regions are filtered and passed to AmpliconArchitect. AmpliconArchitect analyses local genome structure associated with the focal amplification and the hallmarks of the focal amplification classes present are reported by AmpliconClassifier. These outputs can be packaged with AmpliconSuiteAggregator and uploaded to AmpliconRepository.org, where they are placed in a database which can be queried by users. Users can explore individual samples interactively using plots of copy number and genome region data produced with Plotly and IGV. Users may download collections of results from the site for their own custom bioinformatic analyses.

When deploying AA on replicated samples (spatial replicates, temporal replicates, etc.) derived from the same biological origin, AmpliconSuite-pipeline can utilize the multiple replicates to maximize the information derived from AA. Due to the targeted nature of AA’s analysis, if a seed region is present in one sample, but absent in a related sample due to lower copy number, gauging if a related event exists in the seed-absent sample will be difficult. Similarly, if the boundaries of the seeds are not the same due to evolution of the focal amplification between the two replicates, a fair comparison of the two resulting amplicons will be more difficult. To unify the analyzed genome regions across related specimens, we created a wrapper utility called GroupedAnalysisAmpSuite.py which unifies the seed regions for a collection of related samples. This alternate method creates a union of seed BED files across samples prior to invoking the AA step, ensuring that differences in identified focal amplifications between related samples are not confounded by differences in the seed regions.

The seed regions and the BAM file are passed to AA, which establishes focal amplification boundaries and performs joint analysis of copy number and structural variants in the focally amplified regions. While the AA methodology was described originally in Turner et al., 2017^5^ and refined in Deshpande et al., 2019^9^, we provide the following summary of the method to contextualize its role in AmpliconSuite-pipeline. AA constructs a local genome graph comprised of amplified genomic segments and the SVs joining the segments, where both genomic segments and SVs have an assigned copy-number based on their read support (termed a “CN-aware breakpoint graph”). Given the seed regions, AA performs multiple cycles of exploration for additional SVs joining the graph regions to other locations in the genome and recruits those genome intervals to the graph, ultimately forming what we termed an “amplicon.” The amplicon graph contains three general types of SV edges, “concordant edges” that connect directly adjacent segments of the genome, “discordant edges” that connect non-adjacent pieces of the genome (colored by orientation, Supplementary Figure 1), and “source edges” that exit the graph segments to neighboring locations not included in the graph, or which represent SVs with one end located inside the graph and the other end being an unknown location. AA applies a balanced- flow constraint to correct copy-numbers of genomic segments and SV edges, which it solves using convex optimization. AA then explores the genome graph to decompose it into paths and cycles, constrained by the copy-number available. For focal amplifications of simple structure, individual decompositions may capture the full structure of the focal amplification, but in complex cases they typically represent substructures of a larger or more heterogeneous amplicon^21,22^. To improve the quality of results, AmpliconSuite-pipeline automatically detects samples with poorly controlled insert size distribution, where library preparation resulted in a high rate of read pairs marked as discordant (properly paired rate < 90%) and adjusts AA’s parameters accordingly, reducing false-positive SV calls (Supplementary Figure 2).

Outputs from AA are passed to AmpliconClassifier (AC), which applies rule-based criteria derived from SV patterns associated with known focal amplification mechanisms to predict the presence of ecDNA, breakage-fusion-bridge (BFB) cycles using the AA genome graph and decomposed genome paths^10,12^ (Methods – AmpliconClassifier). If neither ecDNA nor BFB are found, AC attempts to more generally categorize the focal amplification as a “complex non-cyclic” focal amplification, characterized by a high degree of structural complexity, or a “linear amplification” characterized by minimal rearrangements^10^. In addition, AC reports summaries of the focal amplification gene contents, copy numbers, and a numerical score for its structural complexity. If ecDNA is detected, AC also applies the ecContext method, described later in this work to distinguish the mechanism of formation of ecDNA.

To provide better control over false-positive focal amplifications, which may arise from unfiltered low complexity and repetitive regions of the reference genome, we created an optional similarity score-based filter in AC (Methods – AmpliconClassifier) which users can deploy when running AC on a batch of biologically unrelated samples. The underlying motivation is that unrelated focal amplifications should not share exact genomic boundaries and multiple identical SVs unless the events are artifactual. Briefly, the filter functions by computing a previously described^12^ similarity score between focal amplifications in two samples using the overlap of focal amplification genome regions and shared SVs. The similarity score is accompanied by a p-value based on a statistical model fit to a distribution of focal amplification similarity scores from pairs of unrelated samples – where the null hypothesis is that the two focal amplifications are from unrelated origins. The similarity scores and associated p-values are used to filter likely false-positive focal amplifications having unexpectedly high similarity across unrelated samples. If the user begins with a collection of unrelated samples (no replicates or multiple samples from the same individual), then the *-- filter_similar* flag can be set when running AC to automate this filtering. If the batch does contain samples from related origins, the user can alternatively run the *feature_similarity.py* script to get similarity scores and associated p-values for all focal amplifications across their samples. Users may then use the similarity scores to filter in a manner that is respective of related samples in their cohort, so as not to filter focal amplifications that have expectedly high similarity scores due to related origins.

### Improvements to AmpliconArchitect

Since AA’s initial release^9^, we have made significant improvements to AA’s SV detection, and reduced the tool’s runtime, particularly on samples with complex focal amplifications. Chief among these improvements was a refactoring of the SV discovery process to remove redundant re-identification of SVs between stages of analysis. A second category of critical improvements involved improvements to the SV discovery process itself. We added an entropy filter to the SV discovery procedure, which reduces SV calls on problematic low-complexity regions of the genome. The filter requires the Shannon entropy of the distribution of nucleotide frequencies in the reference genome sequence on each end of the SV to be at least 0.75, as measured on the sequence of the reference genome matching the positions of discordant reads supporting the SV. Next, we added a second mapping quality filter requiring at least one read on each end of the SV to have a mapping quality of at least 20 (in addition to requiring all SV-supporting reads to have a mapping quality of at least 5). For users who need even more sensitive detection of SVs, we also added AA support for externally provided SV calls. Users may provide AA with an externally generated variant calling format (VCF) file containing SV calls which augments the SVs discovered internally by AA.

Other improvements included improvements to AA’s Sashimi^23^-style plots, automated warnings if inputs are improperly formatted or the collection of seed regions is too long (>500 Mbp), improved merging of nearby focal amplifications into single AA amplicons, changes to AA’s random downsampling to make the process deterministic – ultimately ensuring that all of AA’s outputs are deterministic for each set of inputs. Lastly, more user control for parameters such as the amount of read support required to call an SV, and control over the minimum allowed insert size between read pairs for SV calling.

Reference genomes each have their own characteristics and require different annotations. AmpliconSuite-pipeline introduces reference genome support beyond hg19 & GRCh37 released initially with AA. We assembled additional reference genome annotations into the AmpliconArchitect “data repo”; a collection of reference genomes and custom annotation files required by AA, providing support for GRCh38 and mouse genome mm10. We also created a reference genome build called GRCh38_viral, which contains GRCh38 and common oncoviral sequences (Methods – Data repo construction). GRCh38_viral is used for discovery of human-viral focal amplifications which would otherwise be missed if oncoviral reference sequences were absent during alignment of reads. AmpliconSuite-pipeline allows users to automatically download these reference annotations from the command-line by setting the *--download_repo* argument.

On initial release, AA was compatible only with python2, however we have added python3 support for all tools available in AmpliconSuite, including AA. Furthermore, we have updated the version support for AA’s dependencies. Most notably the MOSEK package used to balance flow in AA’s genome graph, adding compatibility with MOSEK versions 9 and 10. We anticipate these updates also provide compatibility with the next planned release of MOSEK.

To benchmark the improvements in speed resulting from the updated version of AA, we performed a runtime comparison using the low-coverage (median ∼1x) WGS data from 81 cancer cell line samples with focal amplifications originally analyzed by AA^9^ (Supplementary Table 1). We compared the runtime of AmpliconArchitect version 1.3.r2 against the 2019 version 1.0 runtimes (Figure 2a) on the same system (Intel® Xeon® CPU E5-2643 v2 3.50GHz). We observed that the speedups increased with the length of the original runtime (Figure 2b). Across all samples the median improvement was 2.3x. However, for samples which originally took more than two hours, the median speedup was 6.7x.

**Figure 2:**
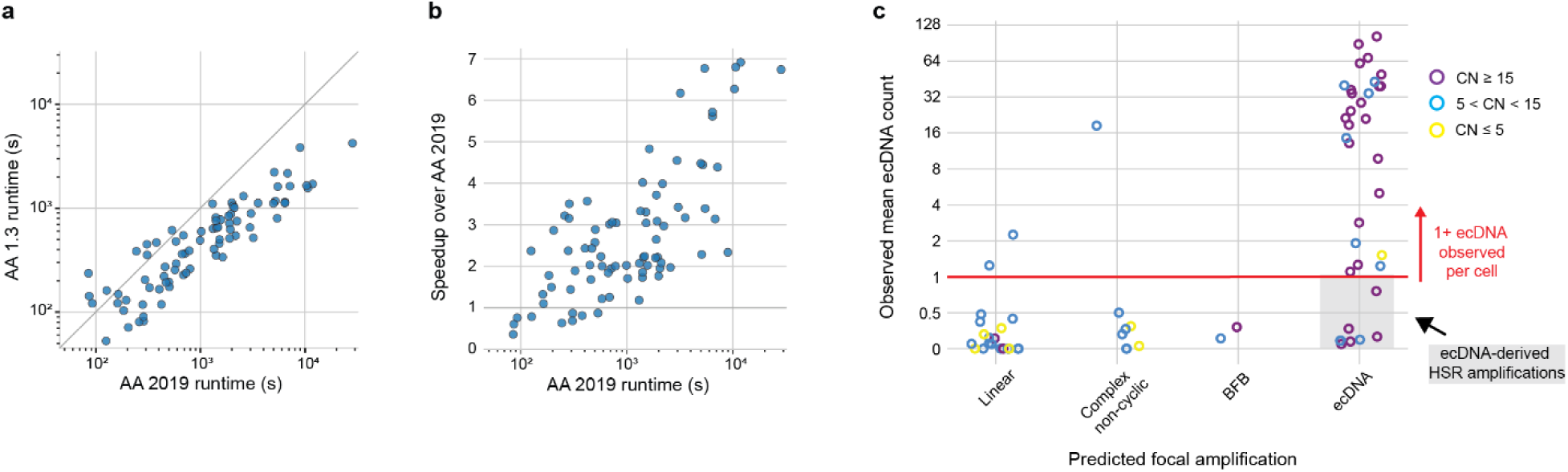
Performance of AmpliconArchitect and AmpliconClassifier. **a)** Runtimes of AmpliconArchitect (AA) on 81 cancer cell line samples using the originally published 2019 version of AA and AA version 1.3.r2. Diagonal line indicates y=x. **b)** Speedup of AA on 81 cancer cell line samples using timing values in panel a. Horizontal grey line at y=1 indicates same speed as original. **c)** AmpliconClassifier predicted focal amplification class for cancer cell line samples with cytogenetically derived ecDNA counts.

To benchmark the ecDNA prediction performance of AmpliconSuite-pipeline, we deployed AmpliconClassifier on a previously described^10^ benchmarking dataset of AA results from 43 cell lines. Across those samples 66 total genes had ecDNA status cytogenetically determined by FISH imaging of metaphase plates, followed by automated image analysis-based counting to establish the number of ecDNA present per cell (Supplementary Table 2). Samples with mean ≥ 1 cytogenetically counted ecDNA per cell were considered ecDNA-positive in the benchmarking dataset. So as not to ignore the critical effects of cancer genome instability and evolution, this benchmarking dataset uses sequencing data gathered concurrently with the cytogenetic data. Applying our updated classification methods (AC version 1.1.1) revealed 90% sensitivity for detection of ecDNA in the dataset (Figure 2c), which was higher than the previous sensitivity (83%) reported in Kim et al., 2020^10^. The three cases with high cytogenetic ecDNA count but ecDNA-negative classification were lacking the necessary SVs to identify an ecDNA-like cycle, illustrating the natural limitations of ecDNA detection imposed by short reads.

### Simplified and flexible installation options

AmpliconSuite-pipeline can be used with or without local installation of the tools. Options not requiring specific installation of AmpliconSuite-pipeline include the GenePattern^16^ module (https://cloud.genepattern.org/) as well as the Nextflow^17^ nf-core/circdna^24^ workflow (https://nf-co.re/circdna). One particularly important benefit of the GenePattern module is that users may upload their data and the analysis will be run entirely by GenePattern. However, we also created containerized images of AmpliconSuite-pipeline for users who may be unable to transmit BAM files outside their local environment and instead wish to deploy a container on their local systems, creating both a Docker and Singularity container for AmpliconSuite. For users who wish to perform standalone installation of AmpliconSuite-pipeline, we created the “AmpliconSuite” package available on Bioconda, and also provided an installer script which can install from source with or without using the AmpliconSuite Bioconda package. Information about all these installation options and how to use them are available in the README file available on GitHub (https://github.com/AmpliconSuite/AmpliconSuite-pipeline). In short, local usage of AmpliconSuite-pipeline involves obtaining the source code & dependencies (via Bioconda, or pulling a containerized image), followed by separately obtaining a copy of the MOSEK license (https://www.mosek.com/license/request), which is free for academic use.

### Interoperability of tools in AmpliconSuite

Tools created to work within AmpliconSuite (AA, AC, CycleViz, ecSimulator, AmpliconReconstructorOM) utilize a common, previously described format^9^ (https://github.com/AmpliconSuite/AmpliconArchitect#file-formats) for representing the genome graph (AA graph format) as well as the genome paths found in that graph (AA cycles format). This common format enables outputs of AA to be utilized by downstream tools such as AC or CycleViz. We also created a conversion script available in the AmpliconSuite-pipeline codebase which allows users to convert the AA cycles and graph file formats into BED files for use with other tools. We created an open-source GitHub organization that encompasses these tools, available at https://github.com/AmpliconSuite.

### Community-sourcing focal amplifications calls with AmpliconRepository.org

To facilitate sharing and exploration of focal amplification calls produced by AmpliconSuite-pipeline, we created a companion website, AmpliconRepository.org (https://www.ampliconrepository.org). AmpliconRepository.org enables browsing of community-provided datasets. Users simply need to register on the site to upload their own AmpliconSuite-pipeline outputs (Supplementary Figure 3). Once a collection of samples is uploaded, the site creates a “project” containing the uploaded samples. Users may opt to make their uploaded project publicly available in the site’s list of public projects. By default, newly created projects will be created privately for the user, and the user can share the private project directly with other users by specifying their email addresses. Users may directly download all results hosted for entire projects, or they may download results for individual samples. AmpliconRepository.org provides a search functionality from the home page, so users can search by focal amplification type or gene name to find results of interest.

Each project page on the site (Supplementary Figure 3) provides interactive graphical overviews summarizing the abundances of focal amplifications present within each sample and across the entire project. The project page also provides a searchable table that lists all samples in the project, including a summary of that specific sample’s focal amplification findings. Previous versions of projects are stored when the user updates a project.

Visiting a sample page (Supplementary Figure 3) provides interactive plots showing the genome-wide copy numbers with locations of focal amplifications highlighted. Users may click on the highlighted focal amplification regions to display AA’s Sashimi-style plot of the CN and SV profile and they may also toggle an igv.js browser^24^ to display the genes in the focal amplified regions. If the project creator provided a JSON file of sample metadata for the sample, that may be displayed or downloaded from the sample page.

As AmpliconRepository projects utilize many different output files from AA and AC, we created a utility called AmpliconSuiteAggregator which allows users to simply provide an archived zip file or tarball file with all their AmpliconSuite-pipeline outputs, and AmpliconSuiteAggregator will perform the necessary reorganization and configuration of the files into the format the site expects. AmpliconSuiteAggregator can upload the files directly to the user’s AmpliconRepository.org account if the user provides their AmpliconRepository username or email address as an argument to AmpliconSuiteAggregator. AmpliconSuiteAggregator is available through GenePattern^16^ (https://cloud.genepattern.org/), or can be deployed on the command-line after installing from GitHub (https://github.com/AmpliconSuite/AmpliconSuiteAggregator).

### AmpliconSuite reveals properties of focal amplifications in large-scale cancer datasets

As a proof of principle of the utility of analysis enabled by AmpliconSuite-pipeline, and to demonstrate its viability for large-scale cancer cohort analysis, we deployed AmpliconSuite-pipeline on 2,525 total combined tumor and cancer cell line WGS samples from the Cancer Cell Line Encyclopedia^25^ (CCLE, 329 cell lines), Pan-cancer Analysis of Whole Genomes^26^ (PCAWG, 2,095 samples), and The Cancer Genome Atlas^27^ (TCGA, 101 samples not present in PCAWG). These collections are hosted as projects on AmpliconRepository.org. We surveyed properties related to the landscape of focal amplification frequency, properties of the focal amplifications themselves, such as their gene contents and copy numbers. We utilized the four classifications of focal amplifications reported by AC. The first two classes are derived from established mechanisms - ecDNA and breakage-fusion-bridge (BFB), and the other two classes reflecting types of homogenously staining region (HSR)-like amplifications without distinct mechanisms of formation, termed “complex non-cyclic” (CNC) and “linear”^10,12^. CNC and linear classes are distinguished by the degree of structural variants in the focally amplified regions^12^.

We found the rate of ecDNA across all analyzed cancer samples was nearly 1 in 4 (24.8%), while the rate of BFBs was approximately 1 in 10 (12.2%) (Figure 3a). We also examined the number of distinct, genomically non-overlapping focal amplifications found per sample and observed 41.6% of all ecDNA+ samples showed coexistence of multiple species of genomically non-overlapping ecDNA, highlighting the diversity of focal amplifications that may exist in a single tumor.

**Figure 3:**
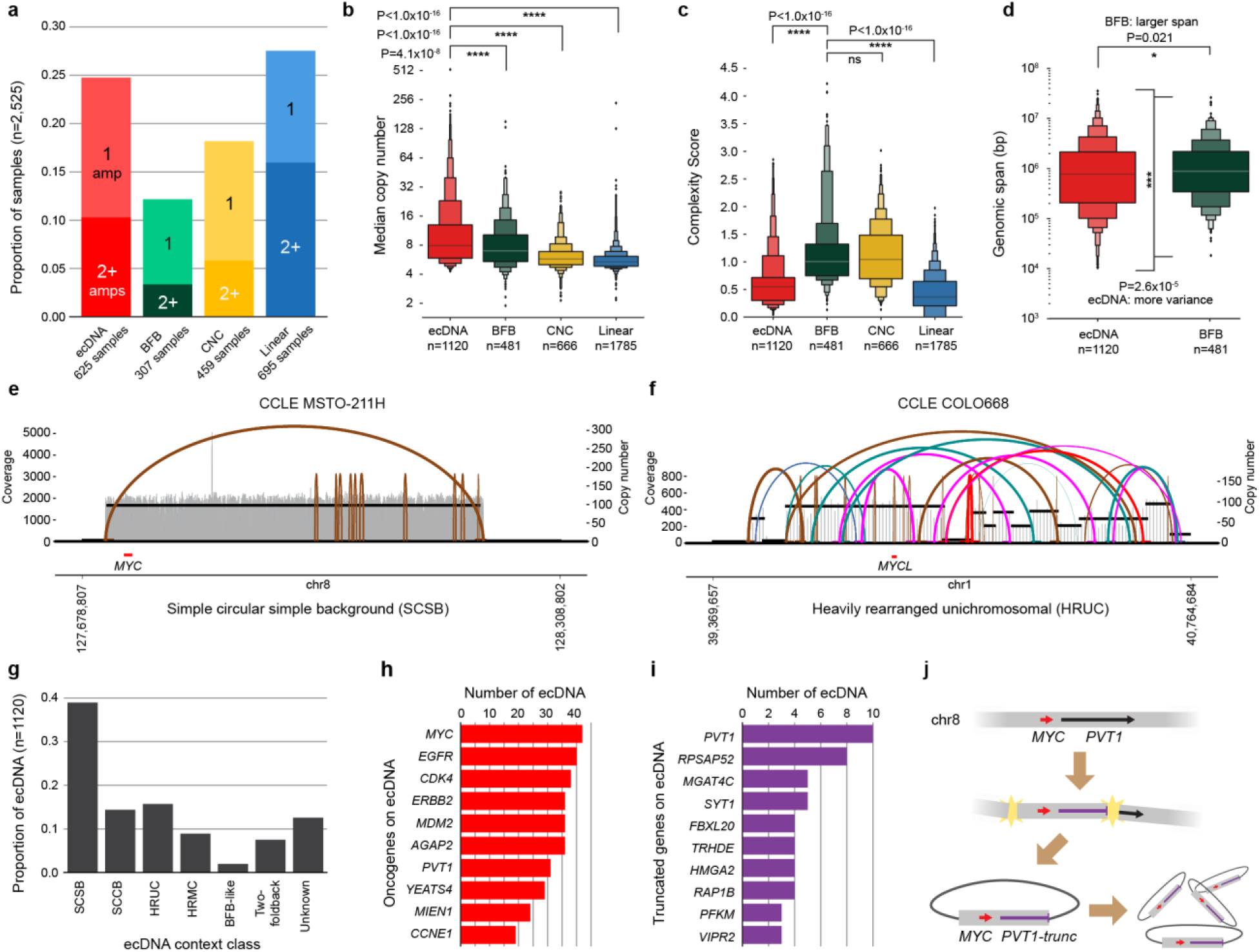
Analysis of focal amplifications in pan-cancer datasets. **a)** ecDNA frequency across combined analysis of CCLE, PCAWG and TCGA samples obtained from AmpliconRepository.org, separated by proportion of samples having only 1, or having 2 or more unique species of that class per sample. **b)** Distributions of the median copy number of each focal amplification, separated by classification, with differences in distribution measured by Mann-Whitney U test. **c)** Distributions of focal amplification complexity scores, separated by classification, with differences in distribution measured by Mann-Whitney U test. **d)** Comparison of distribution of genomic interval span for ecDNA and BFB focal amplifications, with differences in span distribution measured by Mann-Whitney U test and differences in variance between the distributions measured by Bartlett’s test. **e)** An ecDNA predicted by AC having a simple circular simple background categorization, marked by head to tail closure and minimal surrounding rearrangements. **f)** An ecDNA predicted by AC having a heavily rearranged unichromosomal categorization, marked by a high degree of rearrangements involving multiple different genomic regions from a single chromosome. **g)** The frequency of ecDNA context categories across combined CCLE, PCAWG and TCGA samples. SCSB: simple circular simple background, SCCB: simple circular complex background, HRUC: heavily rearranged unichromosomal, HRMC: heavily rearranged multichromosomal. **h)** Frequencies of most recurrently amplified oncogenes on ecDNA. **i)** Frequencies of most recurrently truncated genes (3’ end lost) carried on focal amplifications. **j)** A cartoon illustration of how an ecDNA containing full-length *MYC* and truncated *PVT1* (*PVT1-trunc*) might form, without need for additional rearrangements to later remove the 3’ end of *PVT1* from the amplicon.

To better understand the distinct genomic properties associated with each class, we examined focal amplification copy number, structural complexity, and genomic span. ecDNAs showed significantly higher median copy number (Figure 3b, median CN = 7.9) compared to other classes of focal amplification. BFB showed a median CN of 7.0 (p-value = 4.1×10^-8^), CNC showed a median CN of 5.7 (p-value < 1.0×10^-16^), and linear showed a median CN of 5.4 (p-value < 1.0×10^-16^), with significance evaluated by Mann-Whitney U test. We suggest the significantly higher ecDNA median CN is due to the synergistic combination of non-Mendelian inheritance with positive selective pressure occurring on ecDNA^7^. Despite the higher copy-number, ecDNAs showed significantly lower structural complexity than BFBs (Figure 3c, Mann-Whitney U test, p-value < 1.0×10^-16^), as measured by AC’s amplicon complexity score^12^. This observation is explained by the mechanistically simple way many, but not all ecDNA form, requiring at minimum only two double-strand breaks followed by circularization of the resulting fragment^28^ (Supplementary Figure 4a). BFBs on the other hand require multiple rounds of chromosome arm breakage and end-fusion to achieve high CN^29^, introducing multiple SVs into a single amplicon. We used AC to identify the intervals of the reference genome comprising the focal amplifications and defined the total length of those intervals in each sample as the amplification’s genomic span. BFBs, which are mediated by mechanisms involving whole chromosome arms, tended to have a significantly larger genomic span than ecDNAs (Figure 3d). While we found BFBs had a median span of 880kbp and ecDNA had a median span of 771kbp (Mann-Whitney U test, p-value = 0.021), ecDNAs showed significantly greater variance in their genomic span (Bartlett’s test, p-value = 2.6×10^-5^).

### ecContext categorizes ecDNA by generative mechanism

To understand the contribution of different mechanisms of ecDNA formation to the frequency of ecDNA, we created a method called ecContext for categorizing different types of ecDNA based on patterns of structural variation and copy number, and embedded the ecContext method in AC. Based on previously proposed models of ecDNA formation^28,30–36^ (Supplementary Figure 4), we created seven categories of ecDNA; simple circular simple background (SCSB), simple circular complex background (SCCB), heavily rearranged unichromosomal (HRUC), heavily rearranged multichromosomal (HRMC), BFB-like, two-foldback, and unknown (Methods - ecDNA context analysis, Figure 3e-f, Supplementary Figure 5). ecContext takes as input the AA graph and cycles files, as well as the BED file of ecDNA coordinates in the amplicon produced by AC. It then computes a series of scores which consider the number of copy number states, number of transitions between copy number states, the types of SVs found in the amplicon, and other properties, to place the ecDNA into one of the seven categories (Methods – ecDNA context analysis). We deployed ecContext on the CCLE, PCAWG and TCGA ecDNA amplifications identified with AmpliconSuite-pipeline. We found that SCSB were the most frequent (40.0%), while BFB-linked were the least frequent (1.8%). Together, the two categories most consistent with a chromothriptic origin of ecDNA (HRUC and HRMC) combined for 24.6% of the total ecDNA (Figure 3g). While most ecDNA appear to involve some degree of DNA double-strand breaks (DSBs), we also found that the two-foldback category of ecDNA (7.5%) could be attributed to a proposed model of ecDNA formation which does not involve the use of DSBs, but instead regression of a replication fork. Intriguingly, 12.6% of ecDNA did not conform to any of the mechanistic categories and fell into the “unknown” category, suggesting that there are still unrecognized modes of ecDNA formation. The ecDNA context categories showed differences in the resulting size of the ecDNA, with ecDNA belonging to the simple circular category being the smallest (median 373 kbp), while those in the chromothripsis-associated heavily rearranged categories being the largest (HRUC median 2.046 Mbp, HRMC median 4.927 Mbp, Supplementary Figure 6).

### ecDNA gene amplifications

To illustrate the efficient, positive selection of oncogenes characteristic of ecDNA amplifications, we profiled the density of oncogenes across the various classes of focal amplification. We computed the total number of oncogenes carried by focal amplifications in each class and divided by the combined length of the focal amplifications of the class to compute the oncogene density. ecDNA showed the highest density among focal amplification classes (Supplementary Figure 7, 1664 oncogenes across 2214.6 total megabase pairs), which was more than twice as high as the density of oncogenes found in the hg19 reference genome (997 oncogenes across 3101.8 megabase pairs). Concurring with previous studies^5,10^, we also found recurrent amplification of specific oncogenes across the ecDNAs, with *MYC* and *EGFR* having the highest incidence (Figure 3h). AmpliconClassifier annotates the truncation status of the genes carried on the focal amplifications – detecting if part of the gene was excluded from the focal amplification region. We observed that many genes recurrently showed truncation of the 3’ end when carried on ecDNA (Figure 3i). Some of these identified genes were previously reported to have oncogenic effects when the 3’ end was translocated or lost, including *PVT1*^37^ and *HMGA2*^38,39^. We illustrated how truncated genes could be formed and directly incorporated on ecDNA (Figure 3j), without the need for any prior or subsequent SV operations, ultimately highlighting another mode by which ecDNA enables the formation and amplification of tumorigenic elements in cells.

## DISCUSSION

Detecting focal amplification status from sequencing data reveals critical biology underlying the cancer genome. Cancers with extrachromosomal modes of focal amplification are freed from the constrains of Mendelian inheritance, leading to increased tumor heterogeneity and faster tumor evolution^5,7^. When genome fragments are freed from chromosomes as ecDNA, it is believed they are able to generate multiple new spatial interactions, impacting transcription^1,3,40,41^. To provide a standardized bioinformatic infrastructure in which to study focal amplifications such as ecDNA, we introduced AmpliconSuite. AmpliconSuite contains a workflow called AmpliconSuite-pipeline which simplifies focal amplification detection in paired-end WGS data. At the core of AmpliconSuite-pipeline is the AmpliconArchitect method, to which we have made multiple engineering improvements, and which has improved runtime on complex samples enabling efficient analysis of large-scale cohorts. In this framework, outputs of AmpliconArchitect are summarized by AmpliconClassifier (AC), simplifying the interpretation of focal amplification calls. The utility of AC is also bolstered as we introduced a new method to categorize ecDNA by the mechanism of its genesis.

To benefit the broader community, we built a web platform for community-sourced sharing of focal amplification calls. To this end, we have processed and are hosting focal amplification calls on AmpliconRepository.org from CCLE, PCAWG and TCGA samples. Indeed, other databases of double minute and ecDNA calls exist^42–46^, however they differ from AmpliconRepository.org in many aspects. While most actually incorporate results from AmpliconSuite-pipeline as a source of ecDNA calls, none provide the complete collection of AmpliconSuite-pipeline outputs associated with the samples. More importantly though, many of the existing options do not provide a community-editable platform and lack the ability to create and upload entire projects. By contrast, any user may upload their AmpliconSuite-pipeline outputs to AmpliconRepository by web browser or command-line interface.

By providing a reproducible method to detect focal amplifications and pairing it with the ability to publicly share large collections of focal amplification calls with the community, researchers are empowered to better understand the critical role focal genome amplifications play in driving cancer. Deploying AmpliconSuite-pipeline into environments such as GenePattern and Nextflow provides a critical layer of reproducibility to the package versioning and computing environment, while dramatically improving ease-of-use. Furthermore, the interoperability of tools in AmpliconSuite primes future improvements in ecDNA detection with rapidly maturing long read technologies, as we soon anticipate providing a long-read alternative to AA which could seamlessly fit into this toolset.

We note that there are natural limitations to the detection of focal amplifications, imposed both by sequencing technology and tumor biology. First, we note that AmpliconSuite-pipeline is not itself a sequencing quality-control (QC) package, and users should first perform basic QC checks on their samples before deploying AmpliconSuite-pipeline. AmpliconSuite-pipeline is only designed for analysis of paired-end WGS data, not ATAC-seq, RNA-seq, exome sequencing, or other sequencing strategies. While focal amplifications occurring as subclonal events in a fraction of analyzed tumor cells still exhibit focal amplification signatures in bulk^15^, AmpliconSuite-pipeline is not inherently designed for detection of subclonal events, nor events occurring in highly impure tissue samples. The cytogenetic benchmarking dataset we used was comprised of cancer cell lines, which do not have the same degree of purity- and clonality-related issues as real tumor samples. Furthermore, AA’s copy-number estimates are based on bulk analysis, and superior methods may provide better refinement of the exact copy numbers of these genome regions. Simultaneously, great care should be taken when analyzing genomic copy numbers with other tools to ensure that small (< 1 Mbp) focal amplifications are not smoothed-over due to under-segmentation by CN detection methods which are not designed to detect focal amplifications. Consequently, the best understanding of ecDNA status may come from a combination of sequencing-based methods and cytogenetic approaches to validate the predictions produced by AmpliconSuite-pipeline. Given the instability of cancer genomes, it will be important that cytogenetic data and sequencing data are gathered concurrently from the same biospecimen. Because ecDNA may integrate as HSR or be completely lost from a cancer genome in one single cell division^7^, failure to gather sequencing data and cytogenetics concurrently may lead to inconsistencies.

Our study presented proof of principle analysis using datasets publicly available from AmpliconRepository.org. While the CCLE, PCAWG and TCGA databases represent a wide variety of tumors, the compositional representation of tumors in these cancer databases is not necessarily parallel to other large-scale databases nor representative of population-level cancer incidence rates, and thus the estimate of ecDNA frequency may not precisely represent the true rate of ecDNA in tumors from the global population. Future work will be necessary to completely understand the mechanisms of ecDNA formation. While most ecDNA could be assigned to a context class, some remained unknown - highlighting the need to include additional models of formation to our ecDNA context work. At present, this toolset utilizing paired-end short reads has already proven vital to multiple research groups using the methodology to understand the role focal amplifications play across various cancer types, including medulloblastoma^15^, gastric cancer^47^, esophageal cancer^12,13^, head-and-neck cancer^48^, and more^14,49–51^. Identifying and contextualizing genomic alterations mediated by focal amplifications has the potential to improve therapeutic strategy for patients and speed up progress in the identification of new therapeutic targets that exploit the unique properties of focal amplifications.

## METHODS

### Seed region filtering

Before deploying AmpliconArchitect (AA) to analyze focal amplifications, AmpliconSuite-pipeline filters whole genome copy number variation (CNV) calls to identify candidate regions of focal amplification, called seed regions.

We define the following terminology:

- Copy number segment: A segment, *S* of the reference genome with an assigned copy number *S_CN_*. We denote the genomic length of the segment as |*S*|.
- Chromosome (arm) copy number: The median copy number of genome segments on a specific chromosome (arm). Denoted *C_CN_* or *A_CN_* for chromosome and chromosome arm copy numbers.

The following parameters are used during the seed filtering:

- *CN_cutoff*: The minimum copy number required for a region to be considered significantly amplified, assuming baseline CN=2. The default value is 4.5.
- *Size_cutoff*: The minimum size of a candidate seed region. The default value is 50 kbp.

Seed regions are identified in two phases. First the “pre-filtering” phase which ensures that copy numbers are significantly amplified compared to the chromosome, and secondly on the basis of sequence content and overlap with known problematic elements of the reference genome.

The pre-filtering process begins by taking the CNV calls and computing the median arm CN (*A_CN_*) for each chromosome. The pre-filtering then attempts to remove amplification events consistent with karyotypic abnormalities, as these are not categorized under the definition of focal amplifications. For each CN segment, *S* The first filter checks if the region defined by *S* is longer than 30 Mbp, and if so, discards it (unlikely to be a focal event). Next, if |*S*| is > 20 Mbp, and if so the *CN_cutoff* parameter for the segment is doubled, to prevent small-scale karyotypic abnormalities from being called focal amplifications.

While these filters help handle under-segmented copy-number calls or obvious karyotypic abnormalities, the pre-filtering procedure must also handle CN segments that which fall into long stretches of continuously high copy-number (*S_CN_* > *CN_cutoff* for many consecutive *S*) despite being segmented into multiple different copy number segments. The pre-filter identifies ranges of copy-number segments between 10-20 Mbp in combined length for which *S_CN_ > CN_cutoff* and no two neighboring segments are separated by more than 300kbp of unamplified material. For segments in these ranges *CN_cutoff* is set to 1.5 its default value to ensure the segment is not part of a small-scale karyotypic event.

With these parameters set, the pre-filtering method determines if each segment’s CN (*seg_CN*) passes

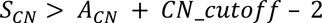

for all copy number segments. Neighboring copy number segments with the same CN are merged. The collection of regions passing these filters we termed the “prefiltered regions”.

AmpliconSuite-pipeline then passes the prefiltered regions to AA’s *amplified_intervals.py* utility^9^, First, the *amplified_intervals.py* script computes coverage statistics from the BAM file to determine *C_CN_*. Computing *C_CN_* from the BAM file independently of the provided CNV calls provides a sanity check. The method checks if

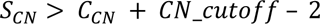

Lastly the seed identification method evaluates properties of the candidate seed regions based on sequence content and overlap with known problem regions of the reference genome using *amplified_intervals.py*. The method filters a candidate seed if it has more than 10% sequence overlapping regions of the genome exhibiting conserved CN gains in normal samples and the candidate also has less than 2 Mbp outside in those conserved CN gain regions (Methods – Data repo construction for details on identification of conserved gain regions). The method then checks the sequence content of the candidate to determine if the mean 35 bp k-mer uniqueness score computed from a sliding window in that region does not exceed 2.5 occurrences in the rest of the genome^9^. For non-oncoviral candidates the method also checks if the candidate is still amplified when accounting for known segmental duplications, checking if

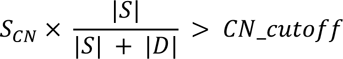

where |*D*| is the total length of the segmental duplication regions overlapping the candidate seed region.

Candidate seeds passing these filters are then merged if they are closer than 300 kbp. Lastly, merged non-oncoviral candidate seeds are filtered based on the *Size_cutoff*, and the final seeds are reported as a BED file with the suffix “AA_CNV_SEEDS.bed”. Users may bypass all these filters and give their own customized seed regions to AA simply by ensuring the CNV BED file provided to AmpliconSuite-pipeline has that suffix on the BED filename.

### AmpliconClassifier

We previously introduced methods to classify focal amplifications from AA outputs in Kim et al., 2020^10^ and later refined the methodology and introduced additional features such as similarity scoring and amplicon complexity in Luebeck et al., 2023^12^. At a high level, AmpliconClassifier (AC) takes the AmpliconArchitect (AA) graph and cycle files describing the CN and SV properties of the AA amplicon and the bioinformatically recovered genome structures present, respectively as inputs and applies a set of rules to predict the type of focal amplification present. The complete description of the rules applied by AC to classify focal amplifications was described in Luebeck et al., 2023^12^, however we provide a summary as follows.

The two inputs to AC are:

1. The AA breakpoint graph file, specifying the structural variant (SV) breakpoint edge locations and the copy numbers of genomic segments in the amplicon.
2. The AA cycles file, specifying a list of cyclic and non-cyclic paths in the breakpoint graph, along with the copy number explained by the path.

first excludes genome paths based on size (<10kbp), overlap with low-complexity or repetitive regions, and those not matching copy number (CN) thresholds (CN of genome regions included in paths must exceed 4.5). Also filtered are paths for which AA’s decomposition assigned a very low copy count. This filter depends on length of path and copy number of genome segments that the path overlaps, but it filters paths with < 3 copies in most cases. For retained paths, AC calculates a length-weighted copy number (*W*) and proceeds with classifications. It initially checks for breakage-fusion-bridge (BFB) cycles, considering the fraction of SV edges (“discordant edges”) with foldback orientation (duplicated inversion with <25 kbp between ends, Supplementary Figure 1), and the fraction of segment pairs in the decomposed paths which are joined by foldbacks. If the proportion of such segment pairs is low (< 0.295) or there are fewer than 3 foldbacks and the fraction of foldbacks across all SVs is < 0.25, or foldback-containing paths represent less than 60% of the total weight, then the amplicon is not considered to contain a BFB. We observed that sequencing artifacts may also take on a foldback orientation, and consequently we added a filter such that a valid BFB must contain no more than 15 foldback edges comprising no more than 80% of the total SVs in the amplicon. If the amplicon does not pass this filter, it is evaluated for presence of complex non-cyclic or linear amplifications. Otherwise, AC marks all paths consistent with BFB as BFB-like paths.

After marking BFB-like paths and cycles (if any), on the remaining paths AC assesses the presence of extrachromosomal DNA (ecDNA), looking for cyclic paths with CN > 4.5, lengths > 50kbp, or at least 12% of the graph’s weight (segment lengths multiplied by CN) being explained by cycles in the genome.

If neither BFB nor ecDNA are found, AC examines paths for complex non-cyclic or linear amplifications. If the fraction of total weight assigned to rearranged non-cyclic paths with (> 5kbp rearranged content) plus the weight assigned to cyclic paths is greater than 0.3 of total *W* across all paths, a complex non-cyclic label is assigned. Similarly, if the ratio of total weight in non-cyclic paths without any rearrangements (no SVs > 5kbp) is greater than 0.25 and there are more than 4 discordant-type SV edges in the amplified region, a complex-non-cyclic label is assigned, otherwise a linear amplification label is assigned. If the cycles are not significantly amplified after applying filters and assessing these checks, a “No amp/Invalid” label is assigned.

Complete resolution of the focal amplification genome structure is often not possible from AA outputs alone due to the structural complexity of an amplification, structural heterogeneity, or the limitations of CN and SV analysis imposed by paired-end short read WGS. However, the signatures of the focal amplification events still exist allowing for resolution of the boundaries of the events. These intervals are extracted from the genome paths and cycles that AC marked in each class with genome segments above default CN threshold 4.5.

AC annotates genes and oncogenes found on the focally amplified intervals using the UCSC RefGene^52^ list packaged in the AA data repo, and oncogene annotations derived from COSMIC^53^ and ONGene^54^ databases.

We have recently introduced methods to remove potential false positive focal amplification calls from collections of samples analyzed from unrelated individuals or biospecimens, based on the assumption that structurally identical focal amplifications should be highly improbable across unrelated samples unless those focal amplifications are artifacts of the reference genome. The filtering method first computes the similarity scores of all genomically overlapping focal amplifications among the samples analyzed in the batch given by the user. Using the amplicon similarity p-value described previously^12^, AC applies a multiple testing correction similar to the Bonferroni correction, where the threshold for identification of statistically significant similar focal amplifications is set to 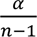 where α is the p-value threshold for significance (default 0.05) and *n* is the number of samples in the collection. All features with a similarity p-value < 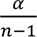 are then filtered. Users can enable the filtering option by setting *--filter_similar* when using AC.

### Analysis of ecDNA context

We developed a rule-based method for ecDNA contextualization, we called “ecContext”, to better understand the frequency of ecDNA origins, including the rate at which ecDNA appear in a chromothriptic context, versus without associated chromothripsis (Supplementary Figure 4). These predictions are produced as a default output by AC. The method takes three inputs:

3. The AmpliconArchitect (AA) breakpoint graph file, specifying the structural variant (SV) breakpoint edge locations and the copy numbers of genomic segments in the amplicon.
4. The AA cycles file, specifying a list of cyclic and non-cyclic paths in the breakpoint graph, along with the copy number explained by the path.
5. The ecDNA intervals file reported by AmpliconClassifier (AC) listing the genomic segments within an AA amplicon which form an ecDNA.

The ecDNA intervals are first expanded by 10kb on both ends to allow for analysis of additional context in the regions surrounding those regions originally marked as ecDNA. ecContext computes six scores based on the ecDNA intervals, considering the segments and breakpoint edges in the graph file that lie completely within the ecDNA intervals (not simply overlapping). To identify the ecDNA-relevant paths in the AA cycles file, ecContext filters out the paths in the AA cycles file which contain genome paths using non-ecDNA SV edges. The remaining genomic segments, SV edges, and paths are then used to calculate the following scores used for context classification:

i. **Cycle fraction** (***C***): This represents the fraction of the length of the ecDNA regions weighted by copy number which can be explained by the heaviest cyclic path reported by AA. First, for each cyclic path comprised solely by ecDNA genome segments, with length *S_P_* and copy number assigned to that path *C_P_*, we calculate its weight as

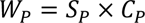

We next define the CN-weighted length of the ecDNA regions, representing the denominator of the cycle fraction. Denoting the sizes of the ecDNA genome segments as *s*_1_, *s*_2_, *s*_3_ … and corresponding copy numbers *c*_1_, *c*_2_, *c*_3_ …, the copy number-weighted length of ecDNA regions, *W_e_*, is calculated as:

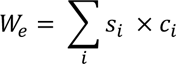

ecContext computes this quantity for each cyclic path comprised solely by ecDNA intervals. For the path *P* with the largest weight, we calculate the cycle fraction as:

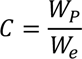

ii. **Number of transitions** (**T**): We defined a transition as a change in copy-number between two genomically adjacent segments in the AA graph in the reference genome. To count the number of transitions in a set of AA graph segments and breakpoint edges, ecContext first filters small (< 5kb) insertions/deletions. The neighboring genomic segments on either side of the filtered indel are joined together if the difference between their copy numbers is less than 2. This updated list of genome segments, having associated copy numbers, is then used to count the number of transitions. If two consecutive genomic segments in the graph are more than 5kb apart, the count of transitions is increased by 1. Alternatively, if two consecutive genomic segments (that are each at least 5kb in size) differ in copy numbers by at least 2, the count of transitions is also increased by 1.
iii. **Copy number states** (***N***): The updated list of merged graph segments computed in the previous step is used to deduce the total number of copy number states in the ecDNA. ecContext takes the copy numbers of the genome segments and sorts them in ascending order. It then iterates through the copy numbers and calculates the average of the group seen so far. Once the next copy number deviates from the current group average by a threshold (default 2), the average copy number of previous group (rounded to the nearest integer) is treated as a copy number state and a new group is started. The total number of groups in the end is the number of copy number states.
iv. **Number of cross edges** (***E***): Two breakpoint edges are considered crossing if the genomic coordinates of one of the breakpoint ends falls between the genomic coordinates of the two breakpoint ends of another edge having both ends on the same chromosome. Any such pair of edges increases the number of cross edges by 1. If any edge connects points from two different chromosomes, it is also considered a cross edge and increases the count by 1.
v. **Number of foldbacks** (***F***): Inverted duplications in the genome give rise to a special class of structural variant called a “foldback” (Supplementary Figure 1). If both mates in a read pair have the same orientation (inversion-like) and the endpoints of the read are mapped no more than 50 kbp apart on the reference genome, we termed it consistent with a foldback SV. We also define the concept of a “foldback fraction” as the number of SV edges that are foldbacks, divided by the total number of SV edges in the ecDNA.
vi. **Unichromosomal or multichromosomal:** We defined a unichromosomal ecDNA as having segments from only one chromosome or having only very small segments (each segment size < 10kbp) from other chromosomes, and a multichromosomal ecDNA as being comprised of significantly large (≥ 10kbp) segments from multiple chromosomes.

Using the six scores computed above, ecContext classifies the ecDNA as one of the following contexts, by checking if they meet the following criteria (examples in Supplementary Figure 5).

The classes are evaluated in the following order and a classification is returned as soon as all criteria of a particular class are met:

1. **BFB-like:** ecDNA may form from a breakage-fusion-bridge (BFB) chromosome^33,34^ (Supplementary Figure 4c). ecContext predicts that the ecDNA is likely to have formed after a breakage fusion bridge event by looking at the number of foldbacks in the ecDNA. If there are at least 4 foldbacks, the fraction of foldbacks among all edges in the ecDNA is > 0.1, the ecDNA is unichromosomal, and has at least 3 copy number states, then ecContext classifies the origin as “BFB-like”. The copy number states must be varied as the copy number profile produced by BFB will be non-uniform in the segments that underwent inverted duplication.
2. **Two-foldback:** ecDNA can form due to regression of a replication fork, following by inappropriate annealing or recombination of the newly replicated strands^35,36^ (Supplementary Figure 4d). This may result in a hairpin-capped double-stranded DNA fragment which is ejected from the fork. Replication of this capped DNA fragment creates a double-stranded circular ecDNA flanked by two foldback inversions. ecContext detects this signature by looking for foldbacks close to (< 50kb) the ends of the ecDNA. If a foldback is found near both ends, and the remaining metrics point to a relatively simple structure – few cross edges (up to 3), low number of transitions compared to number of copy states 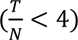, and few copy number states (up to 6) – ecContext classifies the ecDNA origin as “two-foldback”.
3. **Heavily Rearranged:** Chromothripsis gives rise to many ecDNA^31,32^ (Supplementary Figure 4b), and we created the heavily rearranged class to capture ecDNA enriched in a chromothriptic origin. ecContext considers a region to be heavily rearranged if the number of copy number states is high (> 6), or the number of cross edges is high (> 4) or the number of copy-number transitions is high relative to the number of copy number states 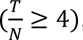, such as would be found in a chromothriptic region with oscillating copy-number^55^. If there is only a single chromosome involved or if the contribution from other chromosomes is minor (< 10kb), ecContext classifies the origin as “heavily rearranged unichromosomal”. Otherwise, the origin is called “heavily rearranged multichromosomal.”
4. **Simple circular**: ecDNA also form from an excisional model mediated by localized double- strand breaks with or without a replication fork being involved, yielding an ecDNA closed by a head-to-tail junction^28,30^ (Supplementary Figure 4a). ecContext checks multiple criteria, described in detail below, to determine if the ecDNA is consistent with a simple circular class, and secondly whether the simple circular ecDNA lies in a genomic background with many genomic rearrangements (complex background), or few genomic rearrangements (simple background). First, if a cycle constituted solely by regions marked as ecDNA by AC cannot be identified in the AA cycles file, then simple circular is ruled out (moving the classification to category 5: unknown). Next, ecContext computes the cycle fraction score (defined in bullet i), including the cycle *P* having the greatest copy-number weighted length. ecContext requires the cycle fraction score (defined in bullet i) be > 0.15 or, in case of an overlapping non-ecDNA linear amplification, it checks if the region defined by the cycle *P* makes up > 30% of the total amplicon length. If these criteria are not met, simple circular context is ruled out. To assess the background genome of the simple circular context, ecContext initially verifies whether cycle *P* (the cycle with the highest CN-weighted length, see bullet i) aligns with an excisional formation model. To determine if *P* represents such a cycle, ecContext checks if *P* is enclosed by a single everted SV edge (Supplementary Figure 1). The presence of ecDNA devoid of extensive surrounding rearrangements, provides compelling evidence for ecDNA formation from an excisional model. Detection of head-to-tail closure involves examining the graph segments constituting *P* to confirm whether the start and end segments (sorted by genome position) define an everted edge on the same chromosome. If ecContext establishes that cycle *P* is enclosed by an everted edge, it records the genome segments used by *P* in a set of genome regions called *R_p_*. In case *P* is not identified as formed by an everted edge, *Rp* remains an empty set. ecContext then computes the previously defined scores ii through vi for three different sets of genome regions. These region sets are defined as follows.
  - Region set *A*: Genome regions marked by AC as the ecDNA, with the 10kbp flanking material added by ecContext.
  - Region set *B*: Region set formed by *A* − *R_p_* . This represents the 10kbp external flanking material outside the original ecDNA regions, and any additional ecDNA regions not captured by *P*.
  - Region set *C*: All regions in the AA amplicon including regions not marked by AC as ecDNA (background or flanking regions), except for the regions in *R_p_*.
  If the scores computed for region set *B* suggest there are few to no external rearrangements – i.e., almost no cross edges (< 2), few copy number states (< 4), few foldbacks (< 4) and unichromosomal content, region set *B*’s scores replace region set *A*’s scores for determination of simple or complex background. Region set *B* replaces *A* to account for cases where there could be a high degree of rearrangement within the simple circular ecDNA, which may occur due to recombination or structural evolution following its formation. Given the scores for each of the three region sets *A*, *B*, *C*, the ecDNA context and background is deduced as follows:
  - Using the region set *A* scores (or *B* scores if replacing *A* with *B*), if the scores point to low complexity – unichromosomal ecDNA, few cross edges (up to 3), low number of transitions compared to number of copy states 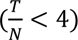, and few copy number states (up to 6), a simple circular context is accepted and the background is deduced.
  - Simple circular context ecDNA are separated into simple and complex backgrounds by considering region set *C* metrics:

o If Region *C* scores point to low complexity (using the same criteria and cutoff values as for *A* or *B*), the final context is “simple circular simple background”.
o Else the final context is “simple circular complex background”.
5. **Unknown:** If none of the above criteria is met, ecContext classifies the ecDNA as having an unknown context. ecDNA falling into this category may include yet heretofore unrecognized modes ecDNA formation, or highly specific modes of formation not captured by broader mechanisms (e.g. two double-strand breaks during a chromosome translocation event subsequently forming multichromosomal ecDNA with otherwise simple circular properties).

### Data repo construction

The AA data repos contain reference genome FASTAs and annotation files used by AA. The hg19 and GRCh37 data repos were previously constructed as described in Deshpande et al., 2019^9^. To construct data repos for GRCh38 (hg38) and mm10 (GRCm38), first we obtained the required reference genome sequences and genome annotation files from publicly available sources. The complete list of files and sources is available in Supplementary Table 3. Broadly, the genome annotation files required in each AA data repo include a BED file of the centromeric coordinates, an excludable regions BED file containing poorly mappable and high-signal regions, a bedgraph file of genome-wide 35 bp mappability scores, a GFF file of genes, a GFF file of oncogenes, a BED file of regions with conserved copy number gains in normal samples, and the reference’s UCSC “SuperDup” segmental duplication annotation file. Lastly, a merged and sorted BED file that combines the excludable regions, conserved gain regions and centromere regions is required by AmpliconSuite-pipeline, named “[genome]_merged_centromeres_conserved_sorted.bed”. While most annotations were available online, we generated 35 bp mappability and conserved gain files as described below.

To create the genome-wide 35bp mappability scores, we used the “gem-mappability” tool (version 1.315) from the GEnomic Multi-tool (GEM) library^56^ to compute genome-wide mappability of 35bp sequences throughout the genome, giving as input the reference FASTA with all non-chromosomal and non-mtDNA sequences removed.

To identify conserved high copy number gains found in normal genomes, we utilized collections of non-cancer genomes. For GRCh38 we used WGS from seven normal samples published in Turner et al., 2017 as well as WGS from HG003, HG006, NA12878 released by Genome in a Bottle (GIAB)^57^. For mm10 we used 40 mouse WGS samples from the Mouse Genomes Project^58^ (Supplementary Table 3). We then utilized CNVkit^20^ (version 0.9.7) to create a flat reference against which CNVkit called CNV profiles. We added regions which showed CN >4.9 in at least 20% of the analyzed samples to the conserved gain region file. For mm10, oncogenic regions were subtracted from the list of conserved gain regions to improve detection of mouse ecDNA covering oncogenes.

To construct the GRCh38_viral data repo, we added additional oncoviral reference genome sequences into the reference genome FASTA file for GRCh38, leaving other files the same. Broadly, the oncoviral sequences are the papilloma and hepatitis viral sequences available from PaVE^59^ (https://pave.niaid.nih.gov) as well as from Jiang, et al., 2012^60^. The complete list of sequences is available in Supplementary Table 1 of Nguyen, et al., 2018^61^.

### Analysis of CCLE, PCAWG and TCGA WGS data

Cancer Cell Line Encyclopedia (CCLE) WGS data was obtained from SRA and the FASTQ files were processed by AmpliconSuite-pipeline (version 0.1032.4), which incorporated BWA^18^ (0.7.17-r1188), CNVkit^20^ (0.9.7), AA^9^ (1.3.r1), however we classified amplicons with a newer version of AC^10,12^ (v1.1.1). TCGA (dbGaP study accession phs000178.v11.p8) and PCAWG WGS files were accessed through the Institute for Systems Biology Cancer Genomics Cloud (ISB-CGC; https://isb-cgc.appspot.com/, ICGC DACO application # DACO-753) and the Amazon Web Services Cloud, respectively. The BAM files were processed by AmpliconSuite-pipeline version 0.1344.2 (CNVkit version 0.9.7, AA version 1.3.r2, AC version 0.4.12). Only the samples with at least 10X sequencing coverage and tumor purity of 0.4 were included in the analysis for detection sensitivity. Sample IDs of the obtained WGS files are available in Supplementary Table 4.

### AmpliconRepository.org framework

The core of AmpliconRepository.org is built as a Django^62^-based web application. User authentication is done using Globus Auth^63^, which supports OAuth2^64^ and OPenID Connect^65^ allowing users to authenticate using single sign-on from many educational institutions and commercial identity providers such as Google. Project and sample metadata are stored in a Mongo-compatible document-based database while the AmpliconArchitect output files and images are stored in the server’s local filesystem or in an Amazon Web Services (AWS) Simple Storage Service (S3) bucket for greater scalability and performance. Interactive sample plots are built using Plotly^66^ and igv.js^24^. The AmpliconRepository.org server is run from a docker container on an AWS EC2 server using AWS S3 for files and AWS DocumentDB as the primary database. It is also possible to deploy private AWS-free AmpliconRepository instances using Docker or Singularity and MongoDB on a local file system where privacy or data confidentiality is required.

AmpliconSuiteAggregator is a tool that takes a collection of AmpliconSuite-pipeline outputs and aggregates them into one compressed file. A user can provide either a zip or tarball file of unstructured AmpliconSuite-pipeline output files, which include the CNV calls used for seeding, the AA and AC outputs. AmpliconSuiteAggregator will decompress the input file and iterate through the provided files to group files for each sample by name. The tool will move them into a pre-determined directory structure and at the same time generate an index file (in the form of a .json file) that contains information of each file and its file path for every sample in the collection.

The tool compresses the resulting directory into a zipped file. The user can specify if they want the resulting zipped file to be directly uploaded onto ampliconrepository.org through an API post. Additionally, users can specify if they want to have the Aggregator re-run AmpliconClassifier, and can also provide a two-column text file which specifies current and desired sample names to perform re-naming of the samples.

### Statistical analysis

The relevant statistical tests are named in the Results section and in figure legends. Statistical tests were performed with SciPy^67^ version 1.11.1. All statistical tests were two-sided unless otherwise stated.

## CODE AVAILABILITY

All code developed for this study is available from the AmpliconSuite GitHub organization: https://github.com/AmpliconSuite.

## DATA AVAILABILITY

No new sequencing data was generated for this study. WGS data for CCLE was downloaded from SRA, PCAWG data was downloaded from ICGC (DACO application # DACO-753) and TCGA data was obtained from dbGaP (phs000178.v11.p8). A list of IDs of analyzed WGS samples is available in Supplementary Table 4. AmpliconSuite-pipeline outputs from samples analyzed in the study are available from AmpliconRepository.org.

## ACKNOWLEDGEMENTS

We thank the many individuals who donated their tissue samples to scientific research, without whom this study could not have taken place. This work was delivered as part of the NCI Informatics Technology for Cancer Research program (V.B., J.M., M.R., J.L., E.H., F.K., T.L., B.D., G.P., D.T., and T. T. were supported by grant U24-CA264379). This work was also delivered as part of the eDyNAmiC team supported by the Cancer Grand Challenges partnership funded by Cancer Research UK (CRUK) (P.S.M., CGCATF-2021/100012; V.B., J.L., B.D., G.P., CGCATF-2021/100025; and R.G.W.V., CGCATF-2021/100016) and the National Cancer Institute (NCI) (P.S.M., OT2CA278688; V.B., J.L., B.D., G.P., OT2CA278635; and R.G.W.V., OT2CA278649). P.S.M. is the eDyNAmiC team lead and co-authors R.G.W.V. and V.B. are members of the eDyNAmiC team. Work supervised by V.B. was also funded by National Institutes of Health (NIH) grant R01-GM114362. Work supervised by J.M. was supported by NIH grants U24-CA264379, U24-CA248457, U24-CA258406. T.L. was supported by NCI grant R50CA243876. P.S.M. was supported by a grant from the National Brain Tumour Society and NIH R01-CA238379. H.K. was supported by the National Research Foundation of Korea (NRF) grant funded by the Korean Ministry of Science and ICT (NRF-2019R1A5A2027340), and NRF-2022M3C1A3092022. S.K. was supported by a Korea Health Industry Development Institute (KHIDI) grant funded by Ministry of Health & Welfare (HI19C1348). We thank all the users of AmpliconSuite who have provided their valuable feedback and testing, particularly Owen Chapman, Imran Noorani, Azhar Khandekar, Chris Bailey, Sihan Wu and Kaiyuan Zhu.

## CONTRIBUTIONS

J.L., M.R., J.M., and V.B. conceptualized this study. J.L., B.D., T.L., and E.H. wrote the manuscript. V.B., J.M., P.S.M, and M.R. supervised the work. J.L. created AmpliconSuite-pipeline. F.K., T.L., E.H., J.L., R.A., G.P., R.K., D.T., and T.T. created AmpliconRepository.org. V.D., J.L., M.A. made updates to AmpliconArchitect. E.H. and T.L. constructed the GenePattern module for AmpliconSuite-pipeline. D.S. and P.B. created the Nextflow integration for AmpliconSuite-pipeline. E.H. and J.L. created AmpliconSuiteAggregator and analyzed the CCLE dataset. S.K., H.K., R.G.W.V, and J.L. analyzed the PCAWG and TCGA datasets. B.D., J.L., and V.B. created the ecContext method. All authors provided feedback on the analyses, edited and approved the manuscript.

## COMPETING INTERESTS

V.B. is a co-founder, consultant, scientific advisory board (SAB) member and has equity interest in Boundless Bio, Inc. (BBI) and Abterra, and the terms of this arrangement have been reviewed and approved by the University of California, San Diego in accordance with its conflict of interest policies. J.L. has previously received compensation for consulting services rendered to BBI, and the terms of this arrangement have been reviewed and approved by the University of California, San Diego in accordance with its conflict of interest policies. P.S.M. is a co-founder, chairs the scientific advisory board (SAB) of and has equity interest in BBI. V.D. is founder and manager of the Bioinformatics Shop, LLC and has equity interest in BBI. R.G.W.V. is a co-founder of Boundless Bio. H.K. has received research funds from JW Pharmaceutical. M.A. is an employee of MOSEK ApS. The other authors declare no competing interests.

## SUPPLEMENTARY FIGURES AND CAPTIONS

**Supplementary Figure 1.**
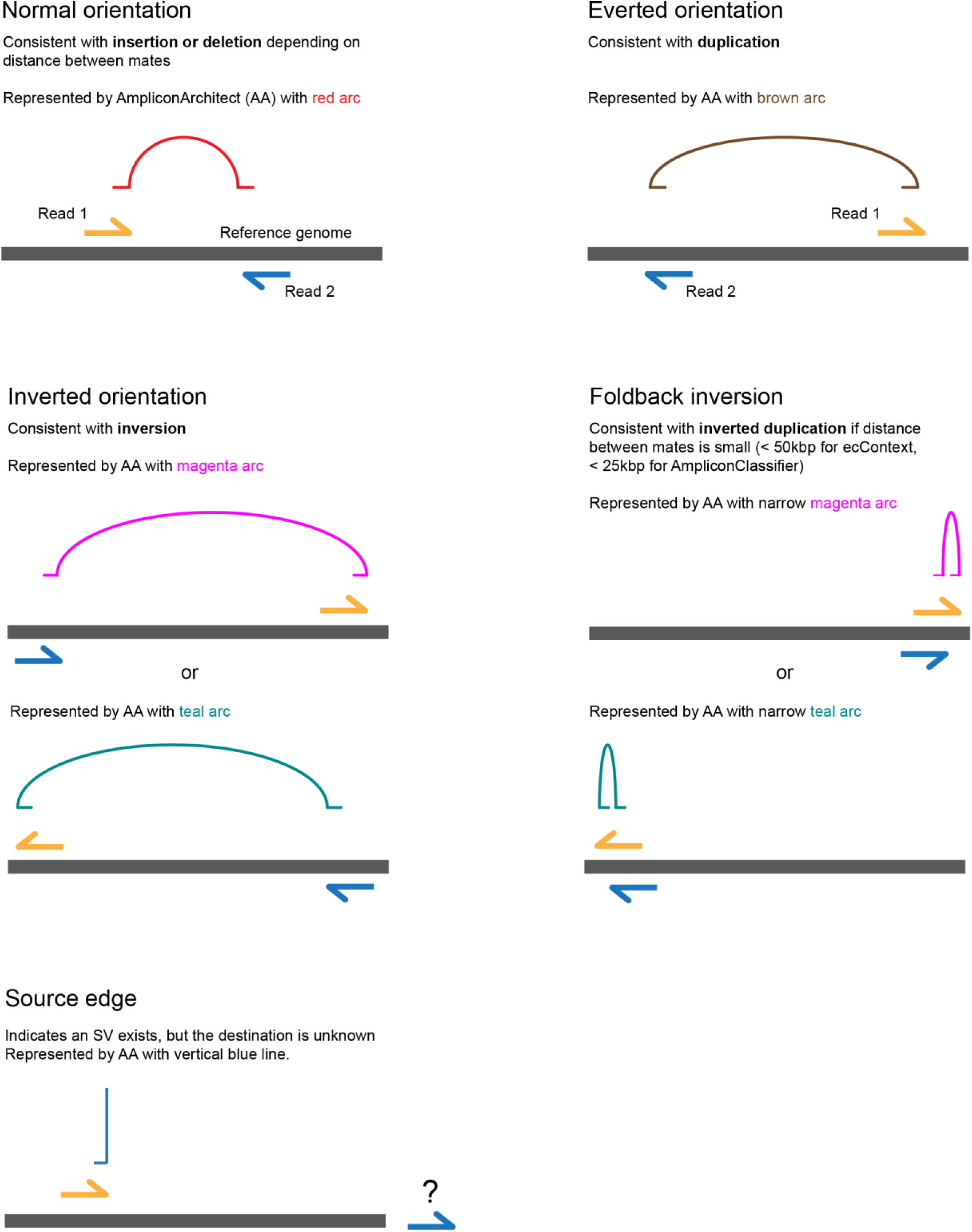
Overview of different types of read pair orientations used by AmpliconArchitect indicating the arc color associated with the SV in its Sashimi-style plots. Reference genome shown has horizontal grey bar with read 1 (orange) and read 2 (blue) indicated. Alignment above the reference bar indicates alignment to the forward strand and alignment below the reference bar indicates alignment to the reverse strand. Inversions may be right-ended (magenta) or left-ended (teal). Foldback SVs represent a type of inversion SV with a distance between mates that is small. Source edges can be comprised of any orientation however one mate in the pair is unmapped or has ambiguous mapping to the reference.

**Supplementary Figure 2.**
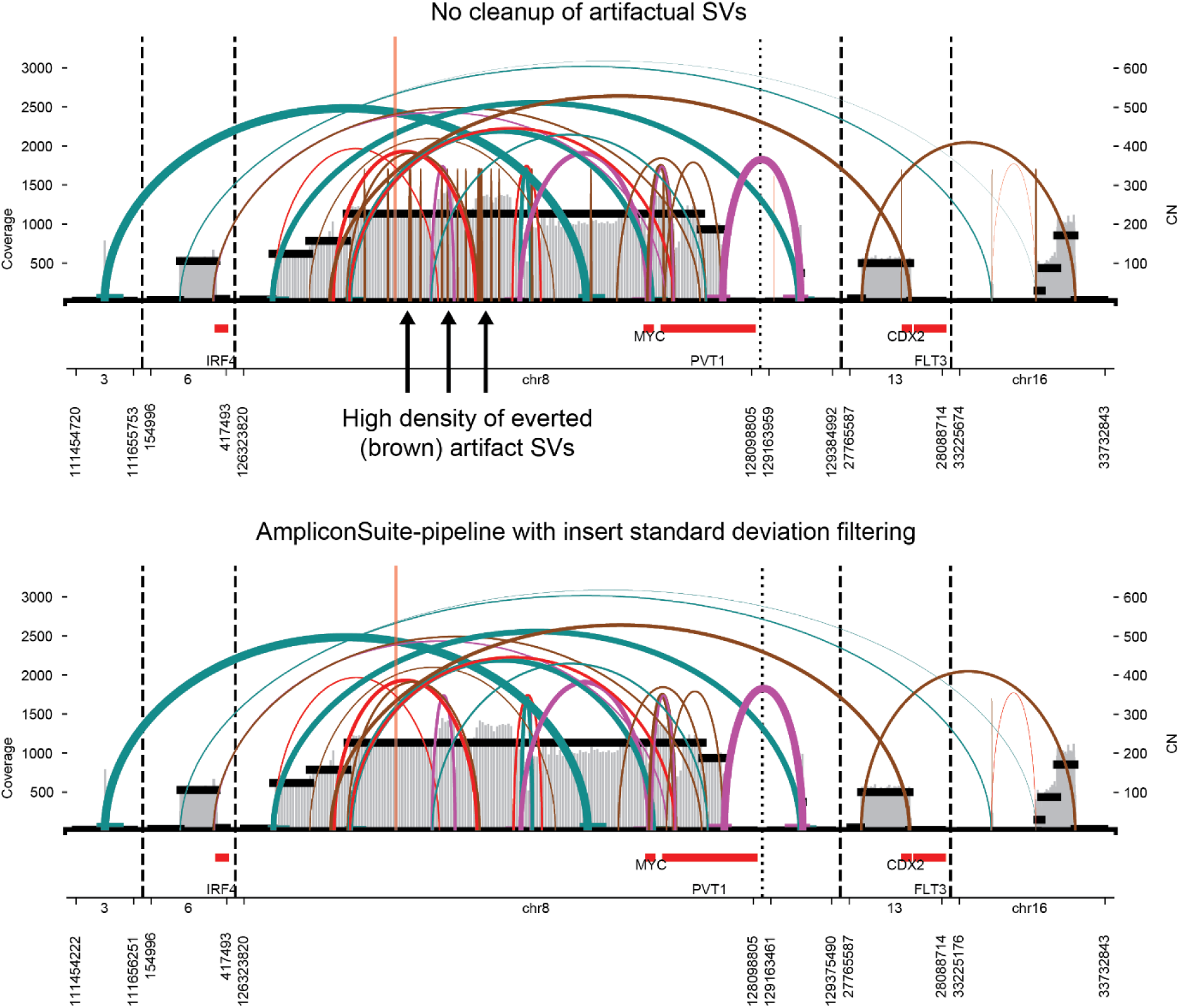
AmpliconArchitect Sashimi-style plots illustrating copy number and identified SVs in the COLO320DM cell line using sequencing data from Wu, et al., 2019. Top panel indicates multiple artifactual SVs driven by a high degree of variance in the insert size distribution which are filtered when using AmpliconSuite-pipeline. Users can gain even more control over this by manually adjusting the *--insert_sdevs* argument. An --*insert_sdevs* value of 9.0 was used in the bottom panel, while the default (3.0) was used in the top.

**Supplementary Figure 3.**
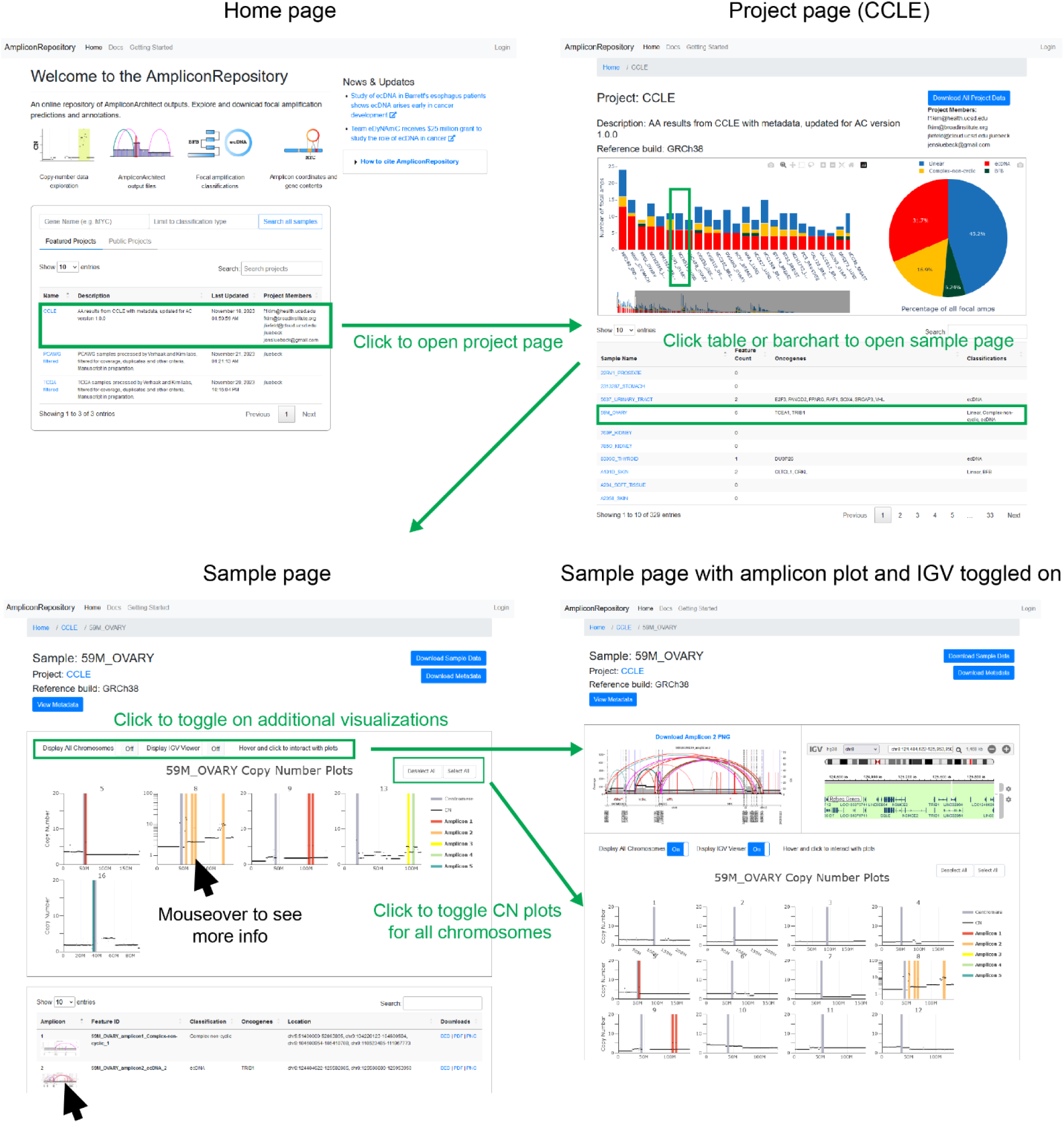
Screenshots AmpliconRepository.org showing how users can select a project by clicking on an entry from the list on the homepage (top left screenshot), which loads a page for that project (top right screenshot). This project page shows Plotly-based visualizations of the number of focal amplifications found in each sample, as well as a pie-chart indicating the overall frequency of each class, and a link to download a tarball of all output files included in the project. Users may click the table listing the samples below or click directly on the bar chart to bring up a page for that specific sample (bottom left screenshot). The sample page shows an interactive Plotly-based visualization of the copy-number data for the sample, and highlighted bars indicating the genomic locations of the focal amplifications. Users may hover or click on elements in the copy number plots to see more information or display the corresponding AmpliconArchitect amplicon image. Users may also hover over the thumbnail image in the table to show a larger image for the amplicon image. The sample page provides links for the user to download the sample data and metadata if provided. Users may toggle on a display of copy number for all chromosomes as well as an IGV display which will render when the user clicks a highlighted bar in the copy number plots (bottom right screenshot).

**Supplementary Figure 4.**
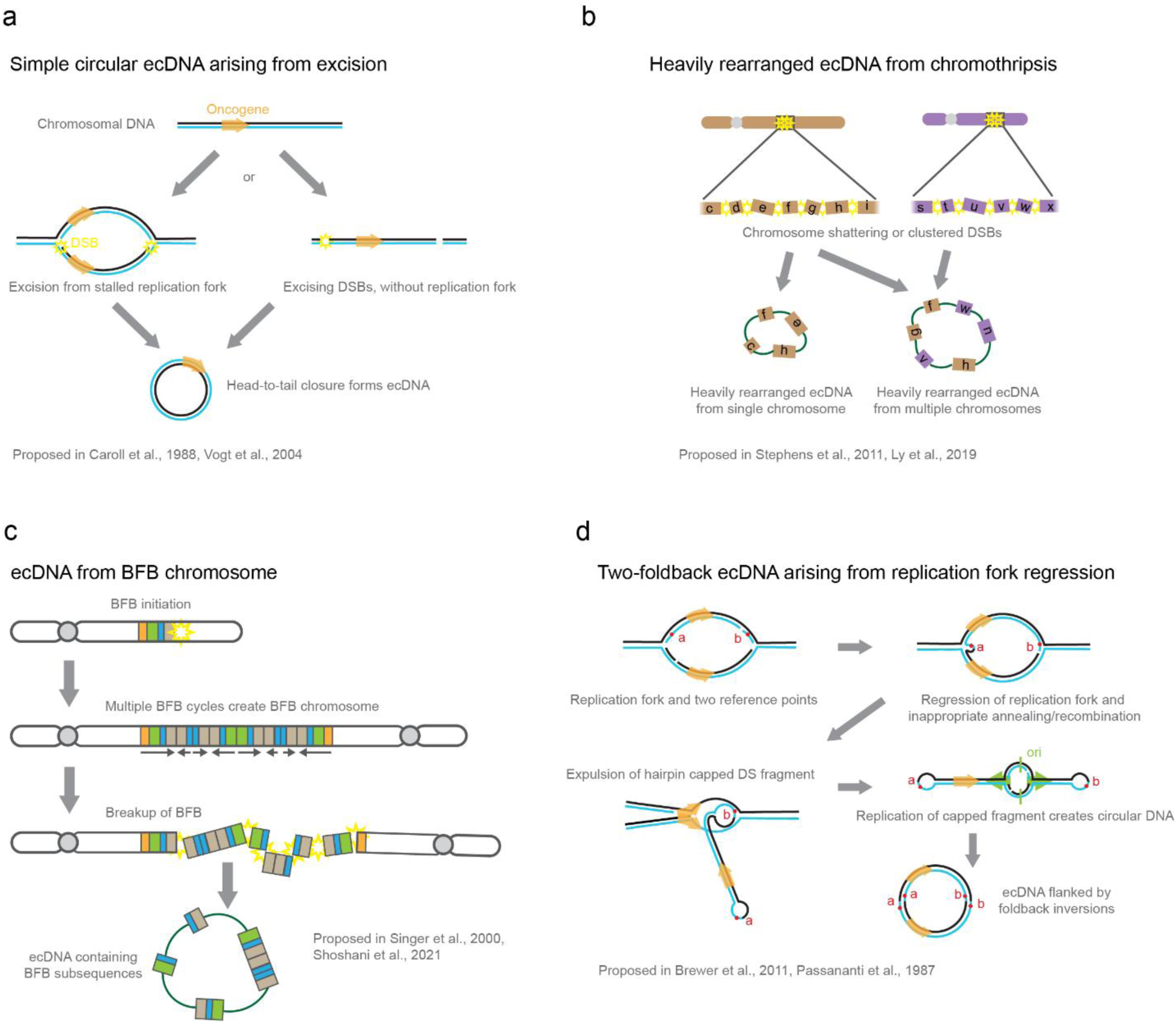
**a)** A model of ecDNA formation based on excision, consistent with models proposed in Carroll et al., 1988 and Vogt et al., 2004. In this model, a fragment of DNA may be excised from a replication fork in crisis, or alternatively may be excised from by two double strand-breaks on pre- or post-replicated DNA. Ligation of the head and tail forms a simple circular ecDNA from that fragment. Additional genomic rearrangements before or after the ecDNA formation are possible. **b)** A model of ecDNA formation which is driven by chromothripsis – the shattering or clustered breakage on genome fragments from one or few chromosomes, as proposed in Stephens et al., 2011 and Ly et al., 2019. While shattering alone does not imply amplification, if the fragments are repaired and joined in a circular fashion, it is possible to generate circular extrachromosomal DNA that includes DNA from one or multiple chromosomes. **c)** A model of ecDNA formation derived from a breakage-fusion-bridge (BFB) chromosome precursor, consistent with observations made by Singer et al., 2000 and Shoshani, et al., 2021. In this model, a BFB chromosome may break apart, either due to chromothripsis or multi-focal breakage of chromatin occurring from physical stress during cell division as the dicentric BFB chromosome is pulled apart. Similarly with the model proposed in **b**, if the fragments are joined in a circular fashion, an ecDNA containing subsequences of the BFB- induced rearrangements is possible. **d)** A model of ecDNA formation based on regression of a replication fork, consistent with models proposed in Brewer et al., 2011 (“ODIRA” model) and Passananti et al., 1987. In this model, a replication fork is shown with two reference points along the newly synthesized DNA of the top strand. Regression of the replication fork followed by either annealing of the handing loose ends of the newly synthesized DNA, or possibly recombination of parts of these strands could lead to a hairpin-capped double-stranded fragment comprised of the two complementary newly synthesized strands. This product would be expelled from the replication fork. Subsequent replication (marked by an origin of replication “ori”) on this fragment would yield a circular double-stranded molecule which contained an inverted repeat of itself, providing the foldback SV bookends we observe in sequencing data.

**Supplementary Figure 5.**
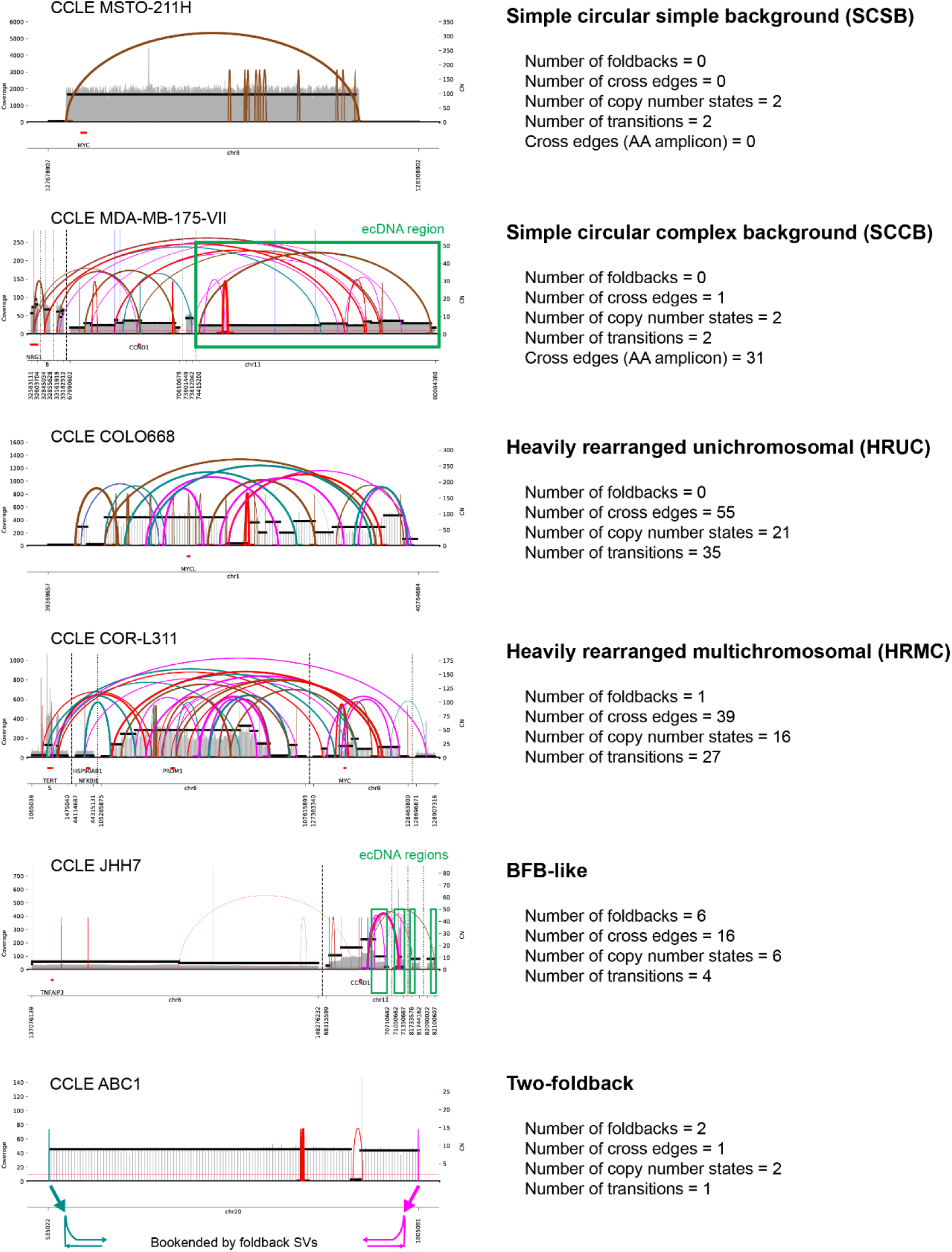
Examples of the six categories of ecDNA context we proposed (not including “unknown”), with values for the scores computed by ecContext which are used when assigning the ecDNA context category.

**Supplementary Figure 6.**
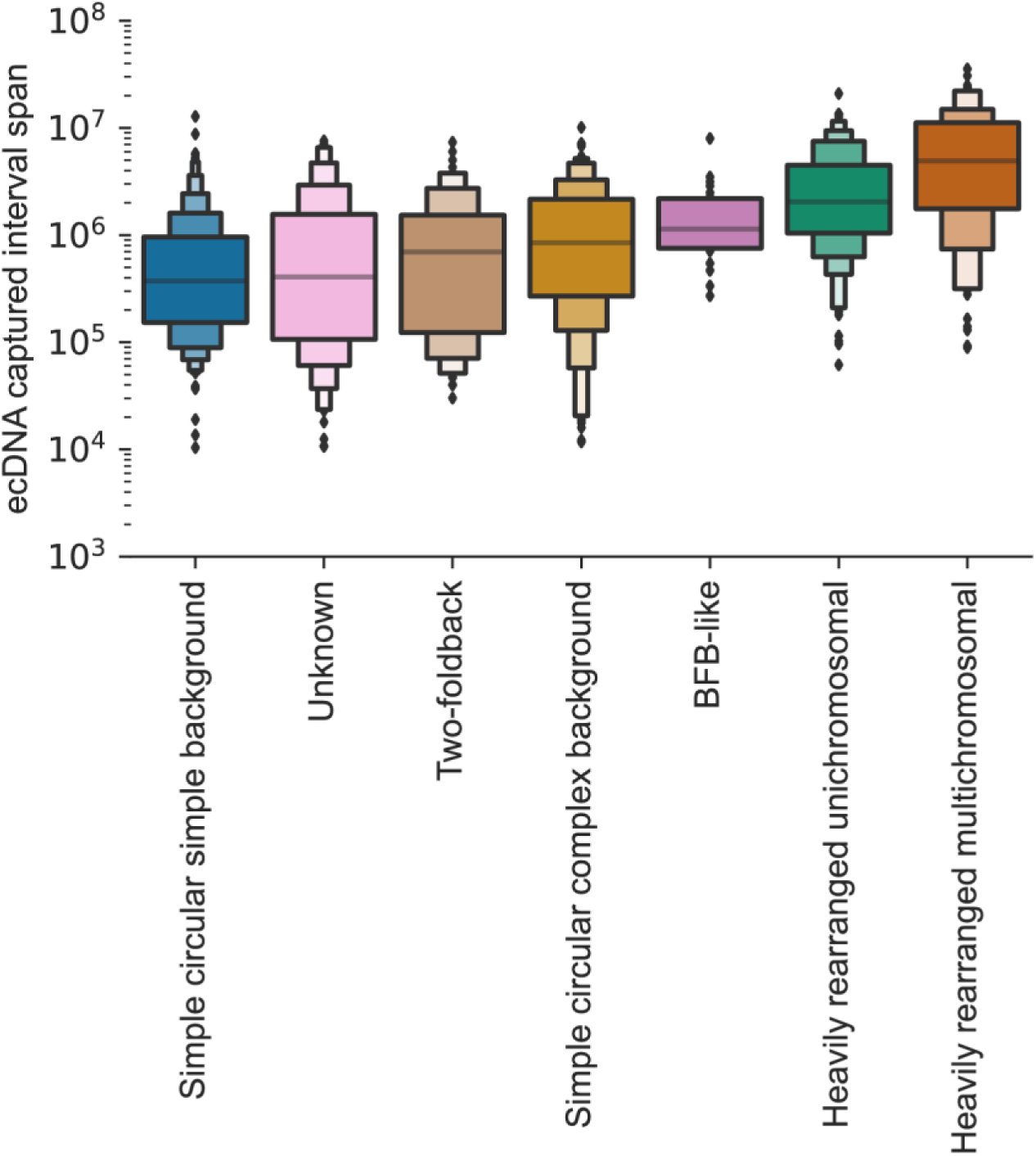
The distribution of captured interval span (length) in basepairs for all ecDNA in each of the categories of ecDNA context reported by ecContext.

**Supplementary Figure 7.**
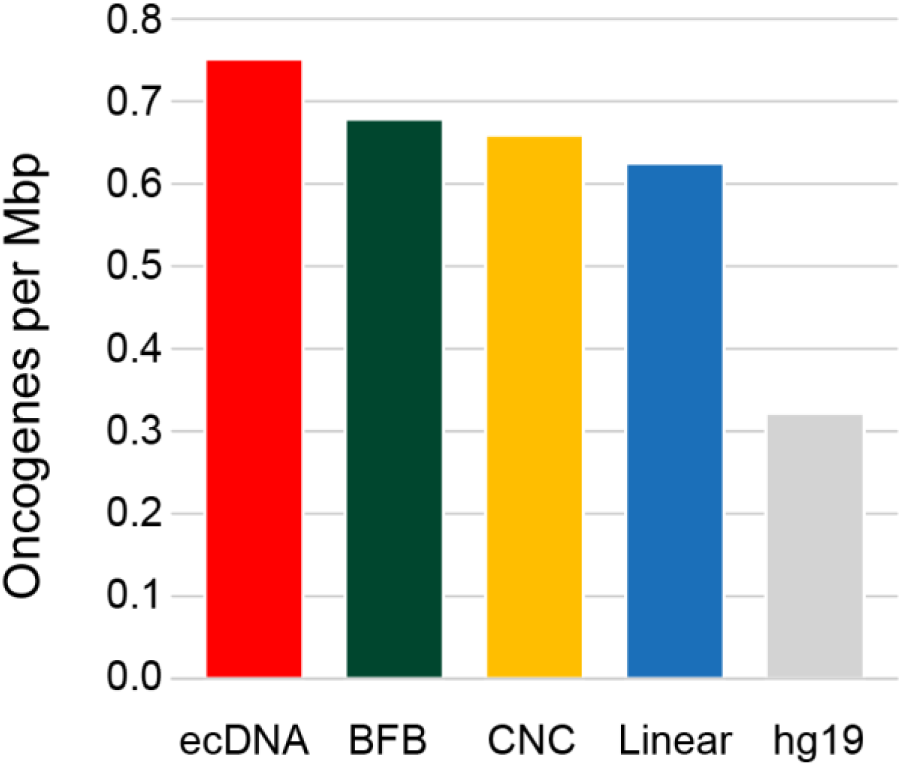
Density of oncogenes across the focal amplification classes using combined focal amplification calls from CCLE, PCAWG and TCGA, with hg19 oncogene density shown for reference.

## Notes

https://ampliconrepository.org/

